# Exposure to relaxing words during sleep promotes slow-wave sleep and subjective sleep quality

**DOI:** 10.1101/2020.12.16.423012

**Authors:** Jonas Beck, Erna Loretz, Björn Rasch

## Abstract

Our thoughts alter our sleep, but the underlying mechanisms are still unknown. We propose that mental processes are active to a greater or lesser extent during sleep and that this degree of activation affects our sleep depth. We examined this notion by activating the concept of “relaxation” during sleep using relaxation-related words in 50 healthy participants. In support of our hypothesis, playing relaxing words during non-rapid eye movement sleep extended the time spent in slow-wave sleep, increased power in the slow-wave activity band after the word cue, and abolished an asymmetrical sleep depth during the word presentation period. On the subjective level, participants reported a higher sleep quality and elevated alertness ratings. Our results support the notion that the activation of mental concepts during sleep can influence sleep depth and provide a basis for interventions using targeted activations to promote sleep depth and sleep quality to foster well-being and health.

---

Sleep, in particular deep sleep, is important for our mental^1^ and physical health^2^ as well as numerous vital functions such as the immune^3^ and cardiovascular systems^4^. The amount of deep sleep, referred to as slow-wave sleep (SWS), depends on prior wakefulness^5^ and declines with age^6^. The depth of sleep is also characterized via slow-wave activity (SWA; EEG power in the 0.5–4.5 Hz band), which has been shown to be a valid marker of global sleep pressure^5^ as well as local sleep need, possibly linked to synaptic downscaling^7–9^.

In addition to neurophysiological mechanisms, cognitive processes can also modulate the depth of sleep. For example, the perception of an unfamiliar sleep environment (known as the “first night effect”) typically reduces the amount of SWS^10^ and decreases SWA specifically in the left brain hemisphere^11^. Similarly, ruminating about a failure experience and bedtime worries about a difficult next day or about an early awakening, negatively affected SWS and SWS latency^12,13^. Conversely, inducing positive thoughts and relaxation using music or hypnotic suggestions increases the amount of SWS and SWA during daytime naps and nighttime sleep^14–16^. However, it is still unknown how cognitive processes which are typically initiated before sleep affect sleep depth, minutes to hours after having fallen asleep.

Here, we propose a mechanism that is based on the activation of embodied concepts during sleep: we assume that the degree of activation of specific cognitive concepts varies during sleep, and that activated concepts are capable of influencing sleep depth depending on their semantic meaning. For example, if the concept of a new and potentially dangerous sleep environment is activated, it remains active during ongoing sleep and thereby decreases sleep depth. Conversely, activation of mental concepts related to “relaxation” and “sleep” are thought to promote sleep depth. This prediction assumes that mental concepts are closely linked to their related bodily functions. According to the theoretical accounts of embodied or grounded cognition, semantic meaning is stored in multimodal (e.g., auditory, visual, motor, somatosensory) neuronal networks^17,18^. For instance, processing concrete words (e.g., arm-related words like ‘catch’) led to cortical activation in the arm motor/premotor region^19^. Similar results were found for processing words related to abstract emotion (e.g., ‘love’) and abstract cognition (e.g., ‘thought’) involving hand and face motor cortices^20,21^. Thus, the processing of words directly activates associated somatosensory brain functions.

The current study provides an initial empirical test for the previously explained mechanism. We predicted that the activation of concepts related to “relaxation” during sleep would increase the amount of deep sleep. To activate concepts of “relaxation” during sleep, we repeatedly presented words related to relaxation during non-rapid eye movement sleep (NREM, sleep stage N2 and SWS). Previous research suggests that our brain can process the meaning of words and activate multimodal representations during sleep^22–25^. In our study, 50 healthy participants slept in the sleep lab for two experimental nights, in a counterbalanced order (see Fig. 1a for an overview of the procedure). In one night, we repeatedly presented relaxing words such as “relax” and “sea” during NREM sleep. In the other experimental night, neutral words were presented (e.g., “produce,” “materials”). Based on the proposed mechanisms explaining how cognitive processes may affect sleep depth, we predicted that the presentation of relaxing words would extend SWS and increase SWA as well as slow-wave density during the time window of word presentation, compared with the control condition. In addition, we hypothesized that event-related responses to the presentation of relaxing words would induce more event-related power in the slow-wave band compared with control words. To control for possible differences in auditory properties between relaxing and control words, a subset of relaxing and control words was also played in reverse during sleep (see Fig. 1a).

**Fig. 1.**
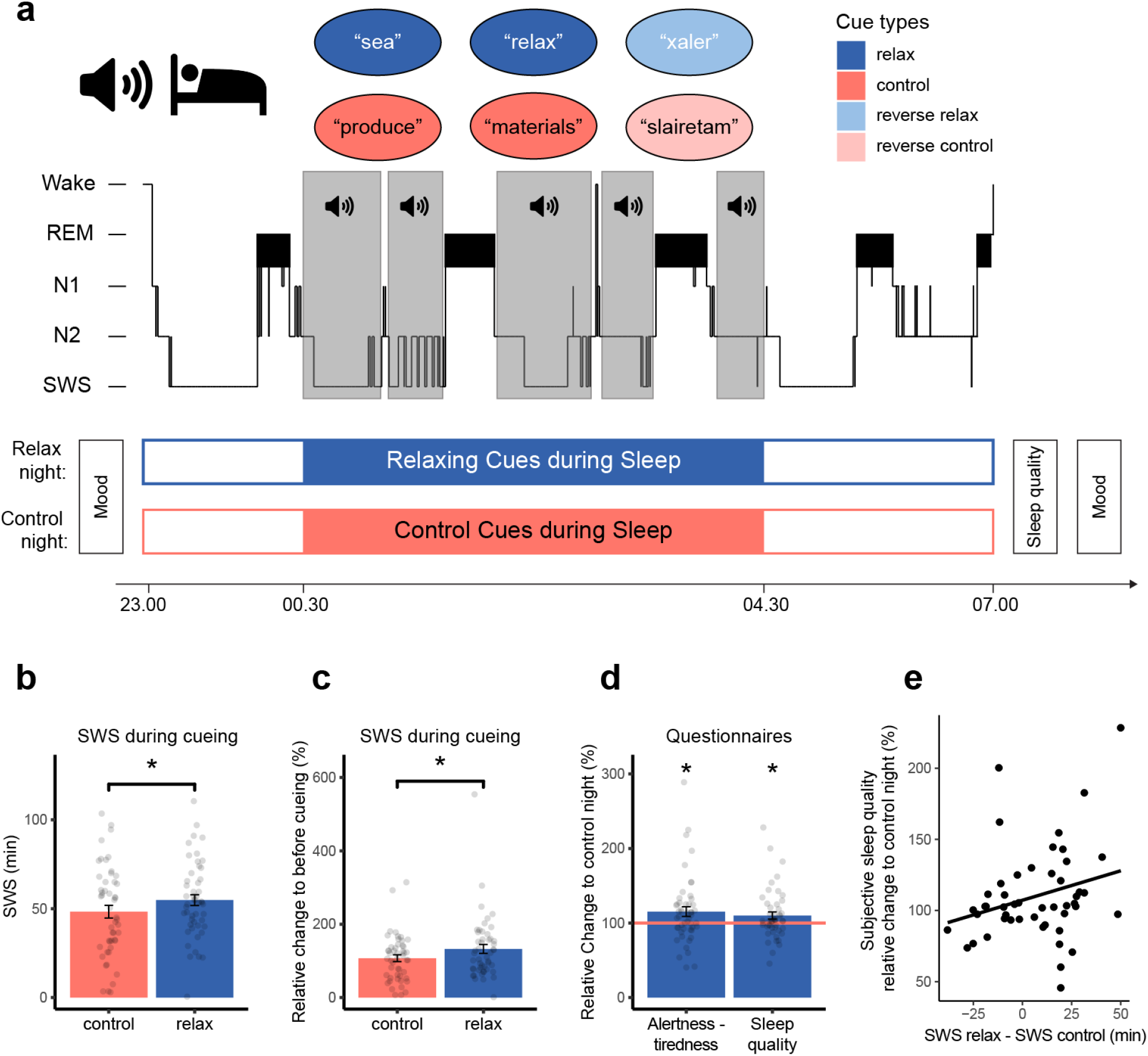
Experimental procedure and sleep results. (**a**) Fifty healthy participants either listened to relaxing words (relax night, blue) or control words (control night, red) during non-rapid eye movement sleep, according to a within-subject cross-over design. During the relax night, forty different relaxing words (e.g., sea, relax) were played in blocks of six words. Each block contained five randomly selected relaxing words (upper circles, blue) and one reverse word (upper circles, light blue). During the control night, the blocks contained five control words (lower circles, red) plus one reverse word (lower circles, light red). When all words were played (i.e., after 8 blocks), the procedure was repeated 15 times, resulting in a total number of 720 word presentations per night. An experimenter monitored the word presentations during sleep and started cueing after the first sleep cycle. Before and after sleep, subjects completed a mood questionnaire. In the morning, a sleep quality questionnaire was conducted. (**b**) Playing relaxing words (blue bar) significantly increased the amount of slow-wave sleep (SWS) in the during- curing period compared with the control night (red bar). (**c**) Playing relaxing words significantly increased the change of SWS sleep from the before- to during-cueing period compared with the control night. One participant was excluded due to high values in the relax night. (**d**) Here provided are the scores in the relax night relative to the control night (set to 100%, red horizontal line). Left bar: Playing relaxing words during sleep significantly increased the change in subjective alertness from before (set to 100%) to after sleep compared with the control night. Right bar: Playing relaxing words during sleep significantly increased subjective sleep quality compared with the control night. One participant was excluded due to missing data. (**e**) The change in SWS from the control to the relax night in the during-cueing period correlated with a statistical trend with the change in subjective sleep quality from the control to the relax night (r_47_ = .26, *P* = .070). Values in the bar graphs are displayed as mean ± SEM. * *P* < .05.

In line with our hypotheses, we found an increased amount of SWS in the period of relaxing word presentation during sleep compared with the period of control word presentation. Furthermore, the increase in SWS induced by relaxing words translated into an increase in subjective sleep quality and alertness reported by the participants. While global SWA and slow-wave density were not altered, we observed a decreased asymmetry of SWA and slow-wave density during the presentation of relaxing words. Finally, event-related processing of relaxing words showed a significantly higher power increase in the SWA band at about 2–3.5 s after stimulus onset.

## Results

After an adaptation night, 50 healthy subjects slept in the sleep laboratory for two experimental nights (8 h) according to a within-subject cross-over design (see Fig. 1a, for an overview of the procedure). Both nights occurred on the same weekday and were spaced one week apart. During one experimental night, relaxing words (e.g., “sea,” “relax”) were presented during NREM sleep (sleep stage N2 and SWS) to promote SWS. During the other experimental night, control words were presented (e.g., “produce,” “materials”). Additionally, relaxing and control words played in reverse were included to control for basic auditory properties of the words (i.e., power spectrum and volume level). Words were presented during NREM sleep starting with the second sleep cycle (at the latest 120 min after sleep onset), as the amount of SWS peaks within the first sleep cycle. Before and after sleep, subjects completed a mood questionnaire. In the morning, a sleep quality questionnaire was conducted.

### Playing relaxing words during sleep promotes SWS and subjective sleep quality

As predicted, playing relaxing words during NREM sleep increased the duration of slow-wave sleep (SWS) compared with the night with control words (see Table 1). This increase was restricted to the time window of word presentation (cueing period). During this cueing period, the duration of SWS was significantly higher in the night with relaxing words (54.81 ± 3.02 min) compared with the night with control words (48.32 ± 3.57 min; *t*(49) = –2.19, *P* = .033, *d* = 0.31; Fig. 1b). The duration of the cueing period did not differ between both nights (172.73 ± 3.71 min vs. 178.68 ± 6.58 min, respectively, *t*(49) = 1.01, *P* > .30). In the before-cueing period (i.e., the first sleep cycle where no words were played), participants showed comparable amounts of SWS (relax: 48.58 ± 2.91 min, control: 47.92 ± 2.25 min, *t*(49) = –0.28, *P* > .70). When controlling for the amount of SWS in the before-cueing period (set to 100%), playing relaxing words increased SWS up to 132.49 ± 12.23%, whereas this increase was significantly lower for control words (107.28 ± 9.03%; *t*(48) = –2.18, *P* = .034, *d* = 0.31; Fig. 1c). One subject had to be excluded in this analysis due to high increases in SWS in the night where relaxing words were played. Descriptively, in the relax condition participants spent less time awake and less time in stages N1, N2 and REM sleep during the cueing period than in the control condition (see Table 1). However, none of these changes in sleep architecture were altered significantly (all *P* > .14).

**Table 1.** Sleep paramters in the before- and during-cueing period. Means in minutes ± SEM. Sleep period time (SPT) including wake after sleep onset (WASO), sleep stage N1 and N2, slow-wave sleep (SWS), rapid eye movement (REM) sleep and sleep efficiency (time asleep / time in bed * 100). **\*** indicates significant differences between control and relax night with *P* < .05

| Parameter | Control night<br>before cueing | Relax night<br>before cueing | Control night<br>during cueing | Relax night<br>during cueing |
| --- | --- | --- | --- | --- |
| SPT | 80.71 ± 2.25 | 79.13 ± 2.41 | 178.68 ± 6.58 | 172.73 ± 3.71 |
| WASO | 1.00 ± 0.30 | 1.22 ± 0.57 | 6.96 ± 1.75 | 4.74 ± 0.93 |
| N1 | 6.84 ± 0.69 | 6.45 ± 0.73 | 11.12 ± 1.71 | 9.01 ± 1.09 |
| N2 | 20.90 ± 1.34 | 19.33 ± 1.32 | 79.63 ± 4.51 | 75.51 ± 3.57 |
| SWS | 47.92 ± 2.36 | 48.58 ± 2.91 | **48.32 ± 3.57** | **54.81 ± 3.02\*** |
| REM | 4.05 ± 0.74 | 3.55 ± 0.69 | 32.65 ± 2.51 | 28.66 ± 1.71 |
| Sleep<br>efficiency | 80.19 ± 2.08 | 81.39 ± 2.03 | 96.87 ± 0.61 | 97.49 ± 0.43 |

The effect size related to playing relaxing words on increasing SWS duration, was in the small to medium range and restricted to the cueing period. The total amount of SWS during the entire night was also comparable between both nights (relax: 136.21 ± 6.68 min vs. control 130.99 ± 7.39 min, *t*(49) = –1.24, *P* > .20, see supplementary Table 1 for sleep architecture across the entire night). However, listening to relaxing words significantly improved subjective sleep quality: participants reported an increased sleep quality the morning after having listened to relaxing words (110.01 ± 4.78%) relative to the night presenting control words (set to 100%; *t*(48) = 2.10, *P* = .041, *d* = 0.30, Fig. 1d). Furthermore, participants reported sleeping significantly deeper (121.53 ± 8.83%), better (113.10 ± 6.05%) and having an ample amount of sleep (118.20 ± 7.07%) in the night with relaxing words compared with the control night (all *P* < .035, all *d* > 0.31, see Table 2). Participants were unable to tell which type of words were presented during sleep. In fact, only one out of all 50 participants reported having heard some words during sleep in one, but not in the other night. Interestingly, the change in subjective sleep quality from the control to the relax night positively correlated with the change in SWS during cueing from the control to the relax night, although the coefficient only reached a statistical trend (*r*(47) = .26, *P* = .070). A significant and positive correlation of these differences was found for the subjective rating of sleeping an ample amount of time (*r*(47) = .32, *P* = .025), while no associations were found between the change in SWS and the change in ratings of sleeping deeper and better (*P* > .70). Finally, participants reported feeling more alert in the morning compared with the prior evening in the relax night relative to the control night (115.31 ± 6.39%; *t*(49) = 2.40, *P* = .020, *d* = 0.34, Fig. 1d, see Table 2 for other scales of the mood questionnaire). On the behavioral level, the presentation of relaxing words during sleep did neither affect vigilance after sleep (as measured by the psychomotor vigilance test) nor memory consolidation during sleep (as measured by a word-pair associate learning task and two verbal fluency tasks, see supplementary Table 2).

**Table 2.** Subjective sleep quality and mood ratings. Sleep quality was assessed with the subscale “sleep quality” and single items from the SF-A/R (Schlaffragebogen A) questionnaire. Here provided are the mean values ± SEM of ratings in the morning from control and relax night. Mood was assessed using the Multidimensional Mood Questionnaire. Given are the morning ratings relative to evening ratings (set to 100%) from the control and relax night. The fourth column displays the change in % of scores in the relax night relative to the control night (set to 100%). Right column displays *P*-values of one sample *t*-tests of these scores against *µ* = 100.

| Parameter | Control night | Relax night | % change with control<br>night set to 100% | *P*-values |
| --- | --- | --- | --- | --- |
| Sleep quality | 3.18 ± 0.12 | 3.29 ± 0.10 | **110.01 ± 4.78\*** | **.041\*** |
| *deep* | 3.27 ± 0.16 | 3.43 ± 0.13 | **121.53 ± 8.83\*** | **.019\*** |
| *good* | 3.43 ± 0.15 | 3.55 ± 0.12 | **113.10 ± 6.05\*** | **.035\*** |
| *ample* | 2.94 ± 0.14 | 3.20 ± 0.16 | **118.20 ± 7.07\*** | **.013\*** |
| *relaxed* | 3.47 ± 0.13 | 3.57 ± 0.11 | 110.27 ± 5.29 | .06 |
| *uniformly* | 2.98 ± 0.15 | 3.04 ± 0.14 | 114.90 ± 8.32 | .08 |
| *undisturbed* | 3.53 ± 0.17 | 3.22 ± 0.15 | 99.90 ± 5.85 | .99 |
| *restless* | 3.55 ± 0.16 | 3.73 ± 0.14 | 105.54 ± 8.60 | .52 |
| Mood (morning / evening) | 96.26 ± 2.53 | 98.13 ± 2.00 | 103.91 ± 2.25 | .09 |
| *good – bad mood* | 93.61 ± 2.48 | 94.32 ± 1.89 | 103.37 ± 2.58 | .20 |
| *alertness – tiredness* | 100.45 ± 5.13 | 106.59 ± 4.64 | **115.31 ± 6.39\*** | **.020\*** |
| *calmness – restlessness* | 98.65 ± 2.45 | 100.07 ± 4.71 | 103.99 ± 3.36 | .24 |

### Playing relaxing words during sleep reduces asymmetry of frontal SWA and slow-wave density

As listening to relaxing words during NREM sleep extended the time spent in SWS, we tested whether relaxing words additionally increased power in the slow-wave activity (SWA) band (0.5–4.5 Hz) during SWS. We analysed SWA during SWS over the frontal lobe in the cueing period and the before-cueing period. We did not find an increase in SWA during SWS by presenting relaxing words (39.53 ± 1.44 µV^2^) compared with control words, in the during-cueing period (39.60 ± 1.65 µV^2^, *t*(49) = 0.04, *P* > .90). SWA during SWS was also comparable between both nights in the before-cueing period (relax: 46.11 ± 2.19 µV^2^, control: 46.15 ± 2.08 µV^2^, *t*(49) = 0.03, *P* > .90). Analysis with the factors condition (relax vs. control), time (before- vs. during-cueing period), and hemisphere (left vs. right) revealed no interaction between time and condition (*F*(1,49) = 0.00, *P* > .90, *η*_*p*_ < .01). A main effect of time reflected the typical decrease in SWA from the first sleep cycle (before-cueing period) to later sleep cycles (during-cueing period; *F*(1,49) = 13.01, *P* = .001, *η*_*p*_ = .21).

In addition, we observed a significant three-way interaction between condition, time and hemisphere (*F*(1,49) = 10.65, *P* = .002, *η*_*p*_ = .18) and an interaction between time and hemisphere (*F*(1,49) = 4.80, *P* = .033, *η*_*p*_ = .09). In the before-cueing period, power in the SWA band was decreased over the left frontal hemisphere, whereas higher values were observed in the right hemisphere. This asymmetric SWA was significant (*t*(49) = –2.23, *P* = .030, *d* = 0.32) and similarly occurred in both nights (Fig. 2a, left panel, before-cueing bars). While playing control words, this asymmetry of SWA remained, with higher SWA in the right compared with the left frontal hemisphere (*t*(49) = –2.08, *P* = .043, *d* = 0.29). However, frontal asymmetry of SWA vanished during the presentation of relaxing words (*t*(49) = 0.25, *P* > .80). Comparing the degree of asymmetric SWA between the before- and during-cueing period in the night with relaxing words, revealed a significant change from a right dominance in the before-cueing period (left minus right hemisphere: –3.11 ± 1.55 µV^2^) to a symmetrical distribution in the during-cueing period (0.37 ± 1.46 µV^2^; *t*(49) = –3.57, *P* < .001, *d* = 0.50; Fig. 2a, right panel, relax bar). No change in asymmetric SWA was observed in the night with control words (*t*(49) = 0.43, *P* > .60). In the during-cueing period, we observed a trend for less asymmetric sleep when participants listened to the relaxing words (0.37 ± 1.46 µV^2^) compared with the night in which control words were presented (−2.59 ± 1.25 µV^2^, *t*(49) = –1.99, *P* = .053, *d* = 0.28).

**Fig. 2.**
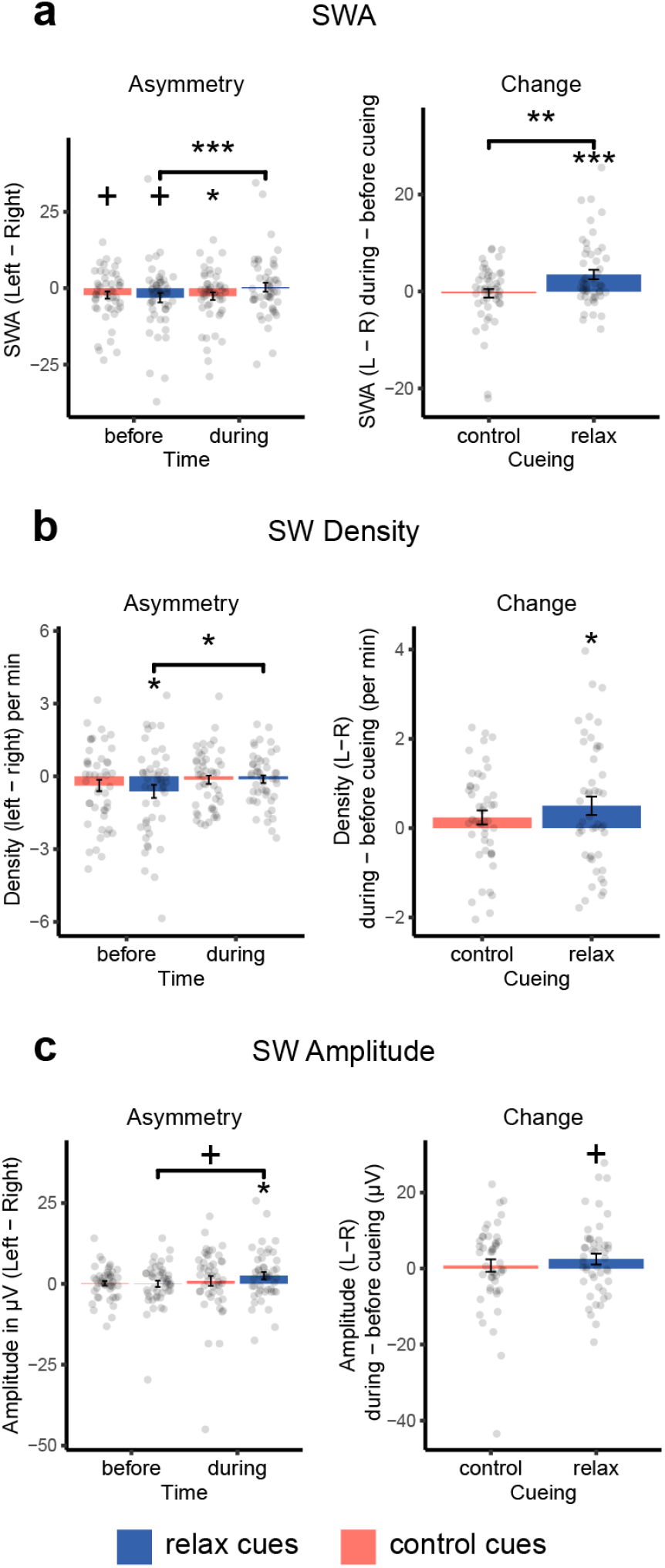
The asymmetry of slow-wave activity (SWA) and slow-waves (SW) during slow-wave sleep (SWS). (**a**) In the before-cueing period, slow wave activity was higher in the right frontal hemisphere compared with the left hemisphere in the relax and control night. The right frontal dominance of SWA persisted during cueing with control cues. In contrast, frontal asymmetry of SWA vanished when relaxing words were played (*t*(49) = -3.57, *P* < .001, *d* = 0.50). (**b**) For SW density, the asymmetric pattern before cueing in the relax night vanished when relaxing words were played during sleep (*t*(46) = -2.42, *P* = .020, *d* = 0.35). The asymmetry of SW density was comparable between the before- and during-cueing period in the control night. (**c**) Asymmetry of SW amplitude (left > right) only occurred during cueing with relax cues (*P* = .033) and increased from the before- to the during-cueing period (*P* = .092). Values are displayed as mean ± SEM. *** *P* < .001, ** *P* < .01, * *P* < .05, ^+^ *P* < .10.

In addition to the general measure of SWA, we analysed single slow-waves detected in the average wave of either the frontal left (Fp1, F3, F7, FC5) or right clusters (Fp2, F4, F8, FC6, see Fig. 1) during SWS. Three participants had to be excluded in this analysis due to an insufficient number of detected slow-waves in the during- or before-cueing period. Consistent with our SWA analysis, we detected a significantly higher density of slow-waves in the before-cueing period over the right frontal cortex (4.55 ± 0.34) compared with the left frontal cortex (4.05 ± 0.26; *t*(46) = –2.25, *P* = .030, *d* = 0.33) in both nights (see Fig. 2b, left panel). When relaxing words were played during the cueing period, the asymmetry of slow-wave density between the left and right hemisphere vanished (left: 2.92 ± 0.21; right: 3.04 ± 0.22; *t*(46) = – 0.75, *P* > .40, *d* = 0.11). The change in asymmetry of frontal slow-wave density, from the before- to the during-cueing period, was significant in the night where relaxing words were presented (*t*(46) = –2.42, *P* = .020, *d* = 0.35, see Fig. 2b, right panel). We observed no significant change in asymmetric slow-wave density in the control night (*P* = 0.13).

When analyzing the negative slope of the detected slow-waves, we also observed steeper negative slopes over the right (–572.20 ± 10.51 µV/s) compared with the left hemisphere (– 530.47 ± 11.79 µV/s; *t*(46) = 4.25, *P* < .001, *d* = 0.62, see Table 3). One additional participant had to be excluded from the analysis of negative slopes due to larger differences between hemispheres in the relax night during cueing. This asymmetry in the negative slope was stable and occurred in the before- and during-cueing period (all *P* < .002). Importantly, no change (before- to during-cueing period) in the asymmetry of the negative slope was observed in both conditions (both *P* > .17), and asymmetry occurred similarly with both relaxing and control words (see Table 3).

**Table 3.**
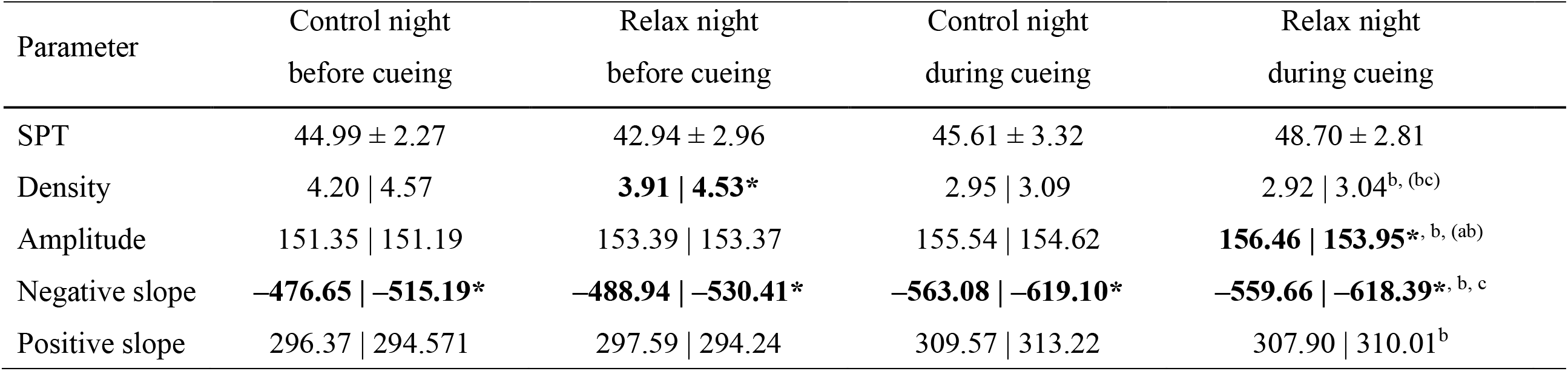
Slow-waves detected in the frontal left or right cluster in SWS. Means of parameters of slow-waves in the frontal left | right hemisphere. Here provided are sleep period time in min (SPT), slow-wave density per minute (Density), peak-to-peak amplitude in µV (Amplitude) and the negative and positive slope in µV/s. (b) indicates a main effect of time (before, during) and (c) a main effect of hemisphere (left, right). (ab) indicates an interaction between cueing and time. (bc) indicates an interaction between time and hemisphere. Bold values marked with * indicate significant differences between the left and right hemisphere. Effects are reported with *P* < .05 (*).

In contrast to slow-wave density and negative slope, the peak-to-peak amplitude of the detected slow-waves was comparable between the right and left hemisphere in the before-cueing period in both nights (*P* > .80) as well as during the presentation of control words (*P* > .50). However, playing relaxing cues during sleep significantly increased the amplitude of slow-waves over the left hemisphere (156.46 ± 4.94 µV) compared with the right (153.95 ± 4.79 µV; *t*(46) = 2.20, *P* = .033, *d* = 0.32, see Fig. 2c, left panel). A statistical trend suggested an increased asymmetry of amplitude in the relax night in the during-cueing period (2.51 ± 1.14 µV) compared with the before-cueing period (0.02 ± 0.97 µV; *t*(46) = –1.72, *P* = .092, *d* = 0.25, see Fig. 2c, right panel). This was driven by an increased amplitude in the left hemisphere in the during-cueing period with relaxing cues (156.46 ± 4.94 µV) compared with the before-cueing period (153.39 ± 5.17 µV; *t*(46) = –2.44, *P* = .019, *d* = 0.36), while the right hemisphere remained on a similar level (*P* > 0.60). Thus, the change in frontal SWA asymmetry observed in the during-cueing period with relaxing words might be explained by differences in slow-wave density and increases in slow-wave amplitude over the left frontal hemisphere together, but not by changes in negative slope of the slow-waves.

### Playing relaxing words during sleep increases event-related SWA

In addition to the effects of relaxing words on general SWS, SWA and slow-waves, we analysed event-related responses to the word presentations during sleep in the time and frequency domain. We compared responses to four relaxing and four control words (each word was presented 15 times resulting in 60 stimuli per category). To control for general auditory properties, the responses were also compared to the exact same words played in reverse.

First, we analysed event-related responses (ERP) of relaxing words (forward vs. reverse) and control words (forward vs. reverse) separately. On the ERP level, forward relaxing words evoked a stronger negative response during three time intervals compared with reverse relaxing words: 770–1306 ms, 1814–2246 ms and 2748–3324 ms after stimulus onset (1^st^ cluster, 24 electrodes, *P* = .006, 2^nd^ cluster: 21 electrodes; *P* = .036, 3^rd^ cluster: 21 electrodes *P* = .040, see Fig. 3a). No significant difference in the ERP responses were observed between forward and reverse control words (see Fig. 3c). When calculating the interaction between condition (relax vs. control) and word type (forward vs. reverse), two clusters in a later period remained significant: A first cluster (*P* = .002) extended from 2662–3346 ms after cue onset, involving 24 electrodes. A second cluster (*P* = .018) extended from 3630–4138 ms after cue onset, involving 22 electrodes (see Fig. 3e).

**Fig. 3.**
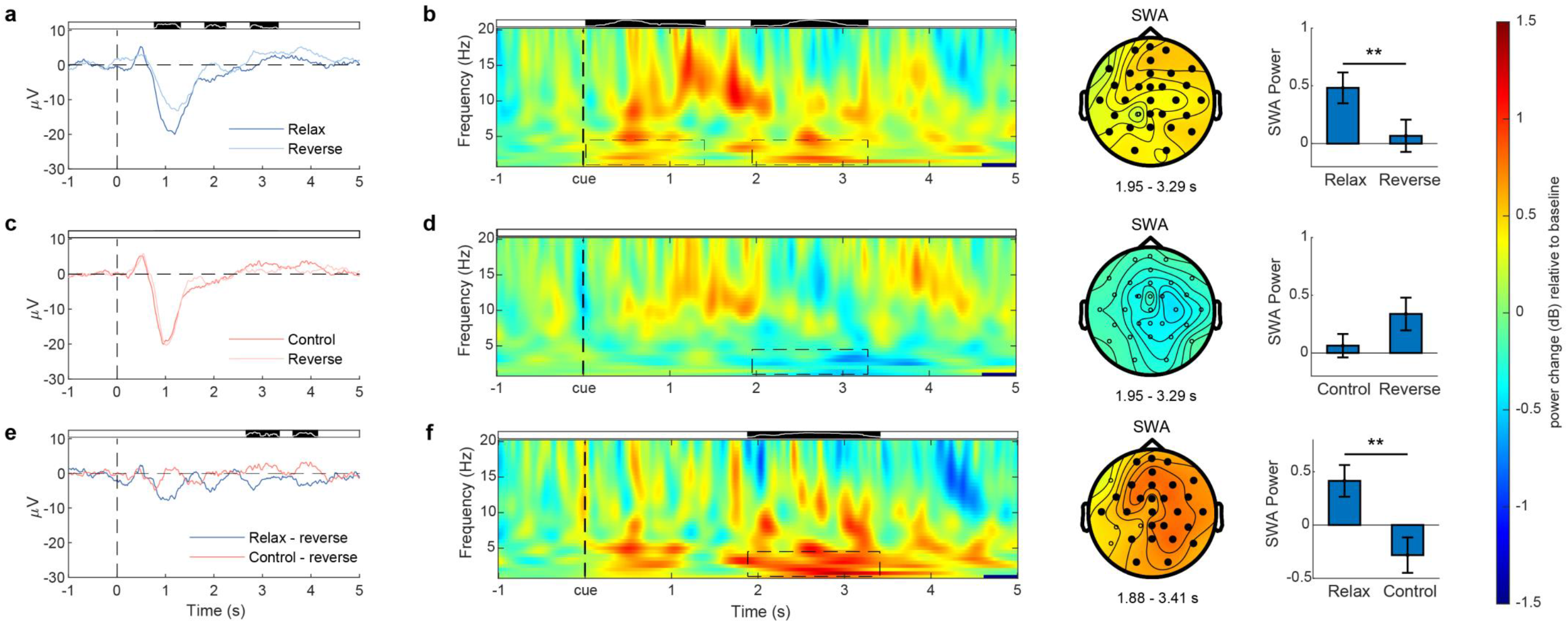
Event-related responses during non-rapid eye movement (NREM) sleep. (**a, b**): Presentation of relaxing words during NREM sleep elicited a higher negative event-related potential than reverse relaxing words (770–1306 ms, 1814–2246 ms and 2748– 3324 ms; **a**) and an increase in SWA power (1^st^: 30–1400 ms and 2^nd^: 1950–3290 ms; **b**, left). The black areas above the plots indicate significant time intervals, while the white line in the black areas displays the number of significant electrodes (full height = 31 electrodes). The increase in SWA power in the 2^nd^ time window was most prominent over frontal and right parietal areas (**b**, middle; significant electrodes = filled black dots). Mean SWA power in the significant 2^nd^ cluster averaged over significant electrodes, duration and frequency band of interest was higher for relaxing vs. reverse relaxing words (**b**, right; *t*(49) = 2.81, *P* = .007, *d* = 0.40). (**c, d**): Event-related potentials (**c**) and SWA power (**d**) of control vs. reverse control words were comparable. Mean SWA power averaged over the same time window (2^nd^) and electrodes as for relaxing words, were comparable between control and reverse control words. (**e, f**): The interaction between condition (relax vs. control) and word type (forward vs. reverse) indicated a higher negative event-related potential for relaxing (– reverse) words compared with control (– reverse) words (2662–3346 ms and 3630–4138 ms, **e**) and an increase in SWA power (1950–3290 ms; **f**, left; see also supplementary Movie 1). The increase in SWA power in the 2^nd^ time window was most prominent over frontal areas and the right hemisphere (**f**, middle, significant electrodes = filled black dots). Mean SWA power in the significant cluster averaged over significant electrodes, duration and frequency band of interest was higher for relaxing (– reverse) vs. control (– reverse) words (**f**, right; *t*(49) = 2.99, *P* = .004, *d* = 0.42). Values in the bar graphs are displayed as mean ± SEM. ** *P* < .01, * *P* < .05.

In the time frequency domain, we specifically focused on changes in the SWA band (0.5– 4.5 Hz). In a separate analysis of relaxing words (forward vs. reverse), higher power in the SWA band was observed in an earlier time window (30–1400 ms, 29 electrodes, *P* = .034, all electrodes except Fp2 and FPz) and a later time window (1950–3290 ms, 30 electrodes, *P* = .034, all except CP1; see Fig. 3b, left panel). The number of concurrent significantly differing electrodes in the second cluster increased until peaking between 2.60–2.65s (30 electrodes), and decreased afterwards (see white line within black bar above Fig. 3b, left panel). Mean SWA power within the second cluster was increased after relaxing words (0.48 ± 0.13) compared with reverse relaxing words (0.07 ± 0.14, *t*(49) = 2.81, *P* = .007, *d* = 0.40, see Fig. 3b, right panel). For the contrast between control words vs. reverse control words, no significant clusters were detected (see Fig. 3d). Mean SWA power, averaged over the time windows and the electrodes of the significant clusters in the relax condition, were comparable between control and reverse control words in the first (*t*(49) = 0.67, *P* > .50) and the second cluster (*t*(49) = –1.66, *P* = .10; see Fig. 3d). When calculating the interaction between condition (relax vs. control) and word type (forward vs. reverse), the cluster in the later period remained significant (1880–3410 ms; *P* = .012; 27 electrodes, all except F7, P7, CP1 and CP5; see Fig. 3f, left panel and supplementary Movie 1). The number of concurrent significantly differing electrodes increased until peaking between 2530–2640 ms (27 electrodes), and decreased afterwards (see white line within black bar above Fig 3f, left panel). Mean SWA power within the cluster was increased after relaxing words (0.41 ± 0.15) compared with control words (–0.28 ± 0.17; *t*(49) = 2.99, *P* = .004, *d* = 0.42, see Fig. 3f, right panel).

## Discussion

In the present study, we present empirical support for our hypothesis that active mental concepts during sleep can influence sleep depth: playing relaxing words during sleep promotes SWS in the cueing period compared with a night in which control words are presented. The increased sleep depth by means of relaxing words was accompanied by a reduced interhemispheric asymmetry of SWA and slow-wave density in the during-cueing period as well as an increase in event-related power in the SWA band several seconds after the cue. The changes observed in objective sleep translated into an increase in subjective sleep quality and alertness ratings.

The findings we reported show that it is possible – in principle – to influence sleep depth by presenting relaxing words during NREM sleep. As an underlying mechanism, we propose that semantic concepts are stored in multimodal representations, and that activation of the semantic meaning will automatically modulate the activation of neural networks responsible for processing the associated bodily function. Activation of the semantic concept of relaxation via the presentation of relaxing words is therefore assumed to activate brain regions responsible for relaxation, sleep induction and maintenance, possibly involving the inhibitory GABAergic system or thalamo-cortical loops, generating slow-waves^26,27^. In addition, activation of meanings related to “relaxation” might also inhibit the activity of arousal- and wake-promoting brain regions in the hypothalamus, brainstem, or basal forebrain^28,29^. Here, we can only speculate on the involved brain regions, as our EEG measures only provide information related to the consequences of the word presentation on the cortical surface.

Still, our event-related analyses revealed distinct increases in power in the SWA band after 2–3.5 seconds, suggesting that the processing of relaxing words during sleep promotes a transient increase in SWA. Importantly, the changes occurred after the typical K-complex-like response to external auditory cues during sleep^30^. Previous research showed that the presentation of single ‘click’ tones in phase with the slow-wave rhythm, enhanced slow-waves during sleep^31^. However, this effect seems to be limited to the presentation of two consecutive ‘clicks’ (ca. 1s), thus lacking enduring effects as well as changes in sleep architecture^32^. Furthermore, the increase in SWA caused by the presentation of relaxing words did not occur for relaxing words played in reverse, thereby controlling for general effects of the tone spectrum, word length and prosody of the auditory stimulus. The increase in SWA induced by relaxing words must therefore be highly specific to the semantic meaning of the word. Importantly, no late event-related increase in SWA occurred when control words were contrasted against control words played in reverse. These findings support our proposition that the activation of semantic concepts related to relaxation, by means of word presentations during sleep, modulates underlying processes which are responsible for slow-wave generation.

The promoting effects of relaxing words on deep sleep were specific to the cueing period and did not yield significant changes of SWS in the entire night. Importantly, a systematically increased sleep pressure or rebound of SWS in the relax night cannot explain this effect, as all sleep parameters were comparable in the before-cueing period. In addition, the effects were specific to SWS, as no other sleep stages were significantly affected by our manipulation. However, the descriptive pattern of decreased amounts of N1, N2, WASO, and REM sleep as well as an increased sleep efficiency during the presentation of relax cues, supports an improved objective sleep quality through relaxing words compared with the control night. Remarkably, participants reported having better sleep quality and elevated alertness in the morning after having listened to relaxing words compared with control words during sleep. These results are controlled for individual differences in sleep perception and general sleep quality, as we applied a within-subject design. Furthermore, the change in the amount of SWS, induced by the relaxing words, positively predicted the improved self-reports of sleep quality. Subjective sleep quality is a highly important marker for the individual evaluation of sleep. As the diagnosis of insomnia is solely based on the subjective experience of the patient and not on objective data^33^, subjective measures are even considered more important for patients and clinicians to diagnose and assess the severity of sleep disturbances^34^. The fact that the presentation of relaxing words during sleep is also capable of improving subjective evaluations of sleep, is therefore highly promising for future applications of such techniques in sleep disturbances like insomnia as well as accompanying affective disorders like depression.

In addition to the effects of SWS and self-reported sleep quality, presentation of relaxing words reduced the typical right-frontal predominance of SWA and slow-wave density during SWS, which was not predicted by our hypothesis. Several previous studies have reported higher frontal SWA in the right compared with the left hemisphere during SWS^11,35,36^. Recently, it was reported that this asymmetry of SWA is even more pronounced in callotosomized patients^37^. The authors argue that such frontal asymmetry of sleep SWA is unlikely to occur due to homeostatic regulation^5^ or functional specification of hemispheres^38^, as extending the time awake leads to a stronger increase of SWA in the left, compared with the right, hemisphere^39,40^.

Thus, it has been speculated that the reduction of SWA in the left hemisphere is related to a “monitoring” or “night watch” system, which watches for potential dangers in the environment, with the ability to induce arousal or wakefulness more rapidly due to a reduced sleep depth^11,22,41^. In support of this notion, hemispheric differences in SWA mainly occur in unfamiliar environments, such as the first night in a sleep laboratory^11,42,43^. In addition, the hemisphere with reduced sleep depth (left) showed an increased evoked brain response to deviant external stimuli while asleep^11^. In line with previous findings, we replicated frontal asymmetry in SWA with reduced SWA in the left, compared with the right, hemisphere in the control night. The same asymmetry occurred in the relax night in the before-cueing period, but vanished when relaxing words were played. A possible explanation is that the activation of the semantic concept of “relaxation” had a “calming” influence on the night watch system, thereby sparing the need of a reduced sleep depth of the left hemisphere. However, this interpretation needs further experimental support.

In a series of previous studies, we have shown that suggestions to relax and to sleep deeper given before sleep, are capable of extending the amount of SWS during a subsequent nap or nighttime sleep^14,15,44^. As a possible explanation we suggested that the mental concept of “relaxation”, given by the instruction to relax before sleep, remains active during subsequent sleep, and is capable of increasing sleep depth by its active multimodal representation^14^. The current study provides direct support for this notion and offers a potential mechanism for the beneficial effects of cognitive interventions given before sleep on later sleep architecture. However, one could have expected that familiarity of the relaxing material before sleep is a necessary prerequisite for the SWS-extending effect of presenting words during sleep. In our study, a subgroup of the participants listened to a tape in which a suggestion to relax and to sleep deeper was given before sleep. The suggestions were identical to the ones used in our previous studies^14,15^. However, we observed no effect of familiarity of the words on the beneficial effects of relaxing words on SWS or other sleep parameters, and therefore decided to combine the data of both groups. We conclude that already existing concepts and associations to the selected words, such as “relaxation” and “sleep”, will similarly activate the multimodal networks related to these concepts, independent of whether these concepts were encountered before sleep or not. To further examine the possibility of targeted memory reactivation of relaxation concepts with verbal cues during sleep, we recommend a learning phase using arbitrary cues, and newly associating them with relaxation-related concepts or experiences (e.g. progressive muscle relaxation or a relaxing virtual reality environment).

A limitation of this study is the choice of words presented during sleep. We presented 40 words from a relaxation text using a metaphor of a fish swimming deeper and deeper into the sea. However, not all of these words might be associated with the same relaxation concept, therefore they might activate other concepts. The associations to one word could also vary interindividually. For instance, “plunge” and “submerge” might even be associated with fear in some people who are afraid of diving for example. Likewise, this should be considered when applying such methods in patients, e.g. with insomnia, where “sleep” might already have negative associations. Moreover, we presented words during NREM sleep, disregarding the phase of the slow-waves. Previous literature suggests that the processing of words is most effective during cortical up states, i.e. at the peaks of slow-waves^45–47^. A closed-loop setup could benefit word processing by targeting the presentation of words precisely in the up-states of slow-waves. This might even strengthen the effect of relaxing word presentations on the depth of sleep. In addition, the volume of word presentations should be individually adjusted to the hearing threshold during sleep. In our study, we kept the volume of word presentations at the same level for all participants and ensured that the average and peak volumes were comparable between conditions. However, the volume of single words varied within conditions (see supplementary Table 4), which likely caused some words not to be processed at all during sleep because they were too silent.

In conclusion, the present study showed that the semantic meaning of words presented during NREM sleep is capable of affecting sleep physiology, SWS maintenance and the subjective evaluation of sleep quality. We argue that the semantic meaning of words presented during sleep is capable of affecting sleep depth by activation of related semantic concepts during sleep. In fact, speaking to people while they are asleep to improve their sleep behavior is a common recommendation in the cases of sleep-walking and night terrors^48,49^, and parents frequently use speech to improve and maintain sleep in their children. Therefore, such presentation of individually-chosen words associated with relaxation and sleep promoting concepts might prove an effective intervention to promote sleep depth and increase subjective sleep quality also in people with sleep disturbances.

## Methods

### Participants

The experiment followed a within-subject cross-over design to compare the effects of the presentation of relaxing or control words during sleep on sleep depth. Fifty-five healthy German-speaking subjects participated in the experiment. Five participants had to be excluded due to insufficient sleep in at least one of the experimental nights (2) or insufficient data quality (3). The final sample consisted of 50 young participants (39 females, mean age = 22.20 ± 3.53 y [M±SD], age range 18–33 y).

None of the subjects were shift workers nor had they been on any intercontinental flights six weeks prior to the experiment. They neither took any sleep influencing medication nor reported any neurological, psychiatric or sleep-related disorders. Subjects confirmed that no surgical interventions had been performed within the three months prior to the experiment, and they did not suffer from impaired hearing. All participants were briefed to wake up at 08.00 h, to refrain from a midday nap as well as to avoid drinking alcohol or caffeine on experimental days. Subjects were paid 130 CHF for participating in all three sessions. The study was approved by the ethics committee of the Canton Vaud (115/15). Participants signed an informed consent after an experimenter explained the study procedure and possible consequences.

### Design and procedure

Subjects participated in three sleep sessions. After an adaptation night, participants slept in the laboratory for two sessions, while polysomnographic data (electroencephalography (EEG), electromyography (EMG) and electrooculography (EOG)) were recorded. Participants arrived between 08.30 p.m. and 09.00 p.m. in the sleep laboratory. Both experimental sessions took place on the same weekday, spaced one week apart and participants filled out questionnaires and performed in memory tasks in both sessions. During one experimental session, relaxing words were presented during non-rapid eye movement sleep (NREM sleep, sleep stage N2 and SWS). During the other experimental session, control words were presented during NREM sleep. The order of condition was balanced across participants according to a within-subject cross-over design. The word presentation protocol was conducted by an experimenter who was blind to the experimental condition. After sleep, psychomotor vigilance, sleep quality and memory recall were assessed.

In addition to the within-subject comparison between relaxing words and control words presented during sleep, the familiarity of the words was varied between subjects. One group of participants (*n* = 37) listened to two texts before sleep. One included the relaxing words, and the other the control words. The participants listened to both texts before both nights. Another group of subjects (*n* = 13) did not listen to the two texts before sleep in any of the two nights. As the effects did not differ between the two groups, we decided to merge the results of both groups.

Subjects completed a paired-associate learning task and two verbal fluency tasks before and after sleep. After waking up, subjects performed a psychomotor vigilance test for 10 min to overcome sleep inertia and assess alertness (see supplementary notes for a detailed description of the three memory tasks and the vigilance test).

### Word presentation during the night

Words were selected and clipped from two texts that we previously used to test the effect of pre-sleep listening on later SWS^14,15,44^, and they are available on our homepage (https://www3.unifr.ch/psycho/en/research/biopsy/). The relaxing text includes a metaphor of a fish swimming deeper and deeper into the sea. It contains many relaxing and reassuring words. The control text comprises a documentation about natural mineral deposits. We selected 40 representative words (see supplementary Table 3, for a complete list of words) from both texts and clipped them from the relaxing text (e.g., to sleep, safe, to relax, fish) and the control text (e.g., surface, deposits, strong, to produce).

As the amount of SWS usually peaks within the first sleep cycle, we started the presentation of the words in the second sleep cycle to ensure sufficient space for a reliable SWS enhancement. An experimenter started to determine sleep stages according to the criteria of the America Academy of Sleep Medicine^50^ with an online sleep scoring setup. Prior to the first experimental night, sleep data of the adaptation night were scanned for each participant to obtain an overview of individual sleep oscillations and architecture. The first sleep cycle was considered complete when a REM episode occurred, and finished, between 60-120 minutes after sleep onset. After the first sleep cycle, words were presented with E-Prime (2.0 SP2, Psychology Software Tools, Pittsburgh, PA) during NREM sleep via loudspeakers placed on a nightstand (distance between speaker and mid of pillow: 85 cm) with an average duration of 1.03 ± 0.27 s (*M ± SD*), and a sound pressure level of 51.34 ± 2.75 dB (see supplementary Table 4 for cue characteristics). If no REM episode was detected, cueing was started no later than 120 min after sleep onset. An experimenter monitored, and manually interrupted, stimulation whenever online polysomnography indicated wake, REM sleep, arousals or movements.

In addition to the main effect of playing cues during sleep on SWS, we aimed to examine event-related responses in the time and frequency domain of the cues. To control for basic auditory properties of words (i.e., power spectrum, volume level etc.), we reversed four relaxing words (easy, to relax, dolphin, sea) and four control words (to produce, magmas, lead, phase). In each night, either relaxing or control words were presented combined with all eight reverse cues leading to a total number of 48 different cues. Cues were presented in blocks of six words containing five randomly selected relaxing or control words and one reversed cue at a random position. This procedure was repeated eight times until all cues had been played once and subsequently started from the beginning. Overall, all cues were presented 15 times in each night resulting in a total number of 720 cues. The presentations of the cues were separated by a random interstimulus interval between 7–9 s. This led to an overall cueing time of 110.3 min in the relax night and 106.2 min in the control night.

### Questionnaires

Subjective sleep quality was measured in the morning with the sleep quality subscale of the SF-A/R^51^. The value of Cronbach’s alpha related to the subscale is .89 in healthy subjects. The scale includes four indices indicating difficulties initiating sleep (1 item), difficulty maintaining sleep (2 items), early awakening with inability to return to sleep (1 item), and general sleep characteristics (6 items). Values between 1–5 indicate if characteristics of good sleep quality are absent (1) or strongly distinct (5). We further analysed the following items (from 1 = not at all to 5 = very much) of the question “*How did you sleep last night?*”: deep, good, ample, relaxed, uniformly, undisturbed and restless. Values are reported as change in % in the relax night relative to the control night (set to 100%).

Mood was assessed before and after sleep using the Multidimensional Mood Questionnaire (MDBF, short form A)^52^. Subjects rated their mood state on 12 items of the question “*In this moment I feel* …” on a five-point Likert scale (1 = not at all, 5 = very much). A total score, and three bipolar mood scales, were calculated (good – bad mood, alertness – tiredness, calmness – restlessness). Cronbach’s alpha of the three scales ranges between 0.78– 0.86, and is 0.92 for the total score^53^. Given are the morning ratings relative to the evening ratings (set to 100%). Similar to the SF-A/R analysis, we analysed the change in % in the relax night relative to the control night (set to 100%).

### Polysomnographic recording

Electroencephalographic (EEG) data were recorded using a 32-channel Easycap Net (Easycap GmbH, Herrsching) with a BrainAmp amplifier (Brain Products, Gilching, Germany), at a sampling rate of 500 Hz, FCz as a physical reference and AF3 as a ground electrode. Two additional electrodes were placed laterally to the outer canthi of the left and right eye to collect electrooculographic (EOG) data. Three bipolar chin electrodes collected electromyogram (EMG) data, and two bipolar electrodes collected electrocardiogram data. Impedances were kept below 10 kΩ for EEG, EOG and EMG electrodes.

For sleep scoring, data were re-referenced against contralateral mastoids, and standard filter settings suggested by the AASM^50^ were applied (e.g., EEG 0.3–35Hz) with an additional notch filter (50 Hz) to reduce noise if necessary. All scorers were blind to the experimental condition. Sleep was scored manually according to the rules provided by the AASM as well as by a central scoring facility^54^. which used a validated scoring algorithm with visual quality control The overall agreement between the two scorings was of 72.97%. A third expert compared both scorings and determined the sleep stage in the case of a disagreement.

### Preprocessing and artifact rejection

EEG data preprocessing was conducted using BrainVision Analyzer software (2.2; Brain Products, Gilching, Germany). Data were filtered using a high- (0.1 Hz) and low-pass (40 Hz) filter with an additional notch filter at 50 Hz and re-referenced to averaged mastoids.

### Power analysis

For the power analysis, data were segmented in 30 s epochs of NREM sleep based on sleep scoring results. Afterwards, data were segmented into equally sized segments of 2048 data points (4s) with 102 points overlapping. Artifacts were rejected automatically^54^ and segments had to pass the following three criteria to be kept: (1) the maximum difference in EMG activity < 150 µV in both EMG channels, (2) maximum voltage step in all EEG channels < 50 µV/ms, (3) maximum difference in EEG activity < 500 µV in all EEG channels. The number of removed segments were manually checked and interpolation was applied if necessary.

We used a fast Fourier transformation (10% Hanning Window, 0.25 Hz resolution) to investigate power differences during sleep. Mean power values (µV^2^) of each channel were exported for SWA (0.5–4.5 Hz) during SWS. Data were further analysed using Rstudio version 1.1.456^55^. Power values exceeding mean activity of all channels by 4 standard deviations were replaced by the mean power separately for each subject. Next, power was averaged over six regions of interest based on topography (see supplementary Fig. 1).

### Slow-wave detection

Slow-wave detection was applied on artifact rejected data from the power analyses in NREM sleep and utilizing a Matlab-based slow-wave-analysis toolbox^56^. Detection followed 4 key stages: first, four reference waves were computed over four regions arranged in a square by calculating the mean activity over 4 electrodes in the same clusters used for power analysis (frontal LR, parietal LR, see supplementary Fig. 1). Second, slow-waves were detected utilizing a local minima approach. As the absolute amplitude of the EEG is influenced by several factors, a relative amplitude criterion (5 standard deviations from the mean negativity) was used to detect local minima. The nearest local maxima served as start and end points of the slow-wave. Only half-waves with a duration between 0.25–1.25 s were kept for further analyses. Each of the 4 reference waves were initially examined independently. Afterwards, potential slow-waves were analysed if a slow-wave was also found within the wavelength of another reference wave. Now, unique slow-waves were found in each reference wave of the four regions. The following parameters were calculated for all detected slow-waves: amplitude (peak-to-peak amplitude in µV), negative slope (between local minima and prior local maxima in µV/s) and positive slope (between local minima and subsequent local maxima in µV/s). In addition, slow-wave density was computed as the number of slow-waves per minute. For further analyses, we focused on slow-waves detected in SWS, which were found only in either the left or the right frontal reference wave.

### Event-related analyses

For analyses of event-related potentials (ERP) and changes in event-related oscillatory power (time-frequency analyses), data were segmented into trials of 14 s based on cue markers starting 7 s before the stimulus onset and followed by the same automatic artifact rejection procedure used in the power analysis. In the night where relaxing words were played, event-related responses to the four relaxing words (easy, to relax, dolphin, sea), as well as their counterparts played in reverse, were extracted. In the night where the control words were played, responses to the four control words (to produce, magmas, lead, phase) and their reverse counterparts were analysed. We used the Fieldtrip toolbox^57^ to compute event-related potentials and time-frequency analysis. Baseline normalization was applied with a baseline period of –1 to 0 s before the stimulus onset. Next, data were averaged per subject and per condition, and grand averages of all conditions were computed. For the time-frequency analysis, a continuous wavelet transformation (complex Morlet wavelets, 5 cycles) was performed on single trials to obtain oscillatory power of frequencies between 0.5–20 Hz in steps of 0.5 Hz and 10 ms.

In the first analysis, we compared relaxing words with reverse relaxing words. Second, we compared control words with reverse control words. Lastly, we computed the interaction by contrasting relax (– reverse relax) with control (– reverse control) words. As we were interested in differences in sleep depth after the cue onset, we focused our analysis on slow-wave activity and averaged over the SWA band (1–4.5 Hz) in the time window 0 to 4.5 s after cue onset across all channels.

### Statistical analysis

Statistical analyses were performed using Rstudio version 1.1.456^55^. Data are presented as means ± standard error. We analysed sleep data (SWS, SWA, slow-waves) using a repeated-measures analysis of variance (ANOVA) containing the within-subject factors cueing (control, relax) and time (before cueing, during cueing). To analyse hemispheric differences, we added the within-subject factor hemisphere (left, right) if applicable. Post-hoc tests for significant interactions and main effects comprised uncorrected paired and unpaired Student’s *t*-tests. Associations were explored using Pearson product-moment correlations. The level of significance was set at *P* < .05.

Results of event-related potentials and time-frequency analysis were compared using cluster-based permutation tests for dependent samples as implemented in the FieldTrip toolbox^57^. The maximum sum of t-values within every cluster served as the cluster-level statistic. Cluster-level alpha was set to .05. To consider the multiple comparisons problem, the cluster-level statistic was calculated for each of 1000 randomly drawn data partitions. The proportion of random partitions exceeding the actually observed test statistic is calculated, resulting in a Monte Carlo *P*-value. The alpha level was set to .05 and corrected for two-sided testing. Alpha level was distributed over both tails by multiplying the probability with a factor of two, prior to thresholding it with the alpha level.

## Acknowledgements

This study was conducted at the University of Fribourg, Department of Psychology, Division of Cognitive Biopsychology and Methods. This work was supported by a grant of the European Research Council (ERC) under the European Union’s Horizon 2020 research and innovation program (grant agreement number 667875). We would like to thank Viviana Leupin for assistance in data collection, Louisa Clarke for helpful comments on an earlier version of this article, and all subjects for their participation.

## Author contributions

J.B. and B.R designed the research; J.B. performed the experiments; J.B. and E.L. analysed the data; J.B. and B.R. interpreted the results, J.B. and B.R. wrote the manuscript.

## Data Availability

The data that support the findings of this study are available from the corresponding author upon reasonable request.

