## Supplementary information for "Exposure to relaxing words during sleep promotes slow-wave sleep and subjective sleep quality"

for

**This file includes:**

Supplementary Materials and Methods
Supplementary Fig. 1. Regions of interest in the power and slow-wave analyses.
Supplementary Fig. 2. Event-related responses of forward and reverse words.
Supplementary Table 1. Sleep paramters in the night with control and relaxing words.
Supplementary Table 2. Task performance in the control and relax cueing night.
Supplementary Table 3. List of words played during non-rapid eye movement sleep (English translation in parentheses).
Supplementary Table 4. Cue characteristics.
Supplementary Movie 1. Event-related slow wave activity power of relaxing compared with control words.

### Supplementary Methods

#### Paired-associate learning task

The word-pair associate learning task (PAL) was conducted[1] and was programmed as well as presented using E-Prime software (2.0 SP2, Psychology Software Tools, Pittsburgh, PA) before and after sleep. In the evening, three blocks were executed: a learning block, a recall block with feedback and a recall block without feedback. Participants learned 80 German semantically associated word pairs (e.g., orchestra – concert) and were asked to memorize as many pairs as possible. Both words of the word pair were presented together for 3000 ms in white font on a black screen. A subsequent black screen was displayed as a random interstimulus interval for 250-750 ms. During the following recall block, the first word was presented, and subjects were asked to type the second word without any time limit. After the input, the correct word pair was displayed for 1000 ms followed by another random interstimulus interval of 250-750 ms (black screen). Afterwards, subjects completed another recall block without feedback. The order of word pairs was randomized for both recall blocks, but the same for all participants. The next morning, another recall block without feedback was conducted with the same word pair order as in the last recall block before sleep. Memory performance was measured as the percentage of correctly recalled word pairs in the last recall block before sleep and in the morning. To assess relative overnight memory improvement, pre-sleep performance was set to 100%.

#### Verbal fluency tasks

During a semantic verbal fluency test, subjects were asked to write down as many words as possible within a given category (fruits, hobbies, profession or animals) in two minutes. The four categories were randomized across the two experimental sessions and the pre- and postsleep session. Multiple named words and words containing the same word stem were excluded. Retrieval performance of long-term memory storage was measured as the number of valid, listed words[2]. To assess relative overnight memory improvement, pre-sleep performance was set to 100%. Comparably, the Regensburger Wortflüssigkeitstest[3] was executed to provide the initial letters (T, N, I, R) of the words to write instead of categories.

**Psychomotor vigilance test**

After waking up, subjects performed a psychomotor vigilance test (PVT) for 10 min to overcome sleep inertia and to assess alertness. Five red zeros were presented as a millisecond counter in the center of a black screen. Participants were asked to press the space bar as quickly as possible when the clock started to run. After the input, the reaction time was displayed in ms for 1 s. As previously suggested[4], the reciprocal response time (mean 1 / RT) was used to assess alertness sensitive to sleep deficits.

**Supplementary figures**

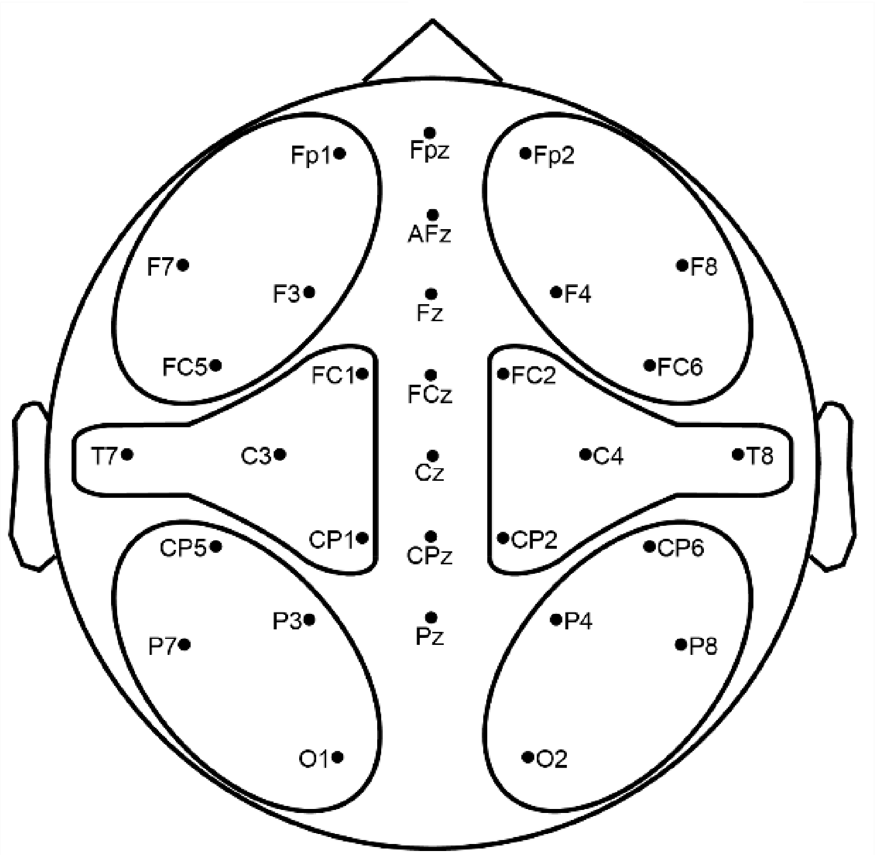


**Supplementary Fig. 1. Regions of interest in the power and slow-wave analyses.** Each cluster comprised 4 electrodes. Power analysis was conducted within all 6 clusters. Slow-wave detection was conducted on the average signal of the frontal and parietal clusters.

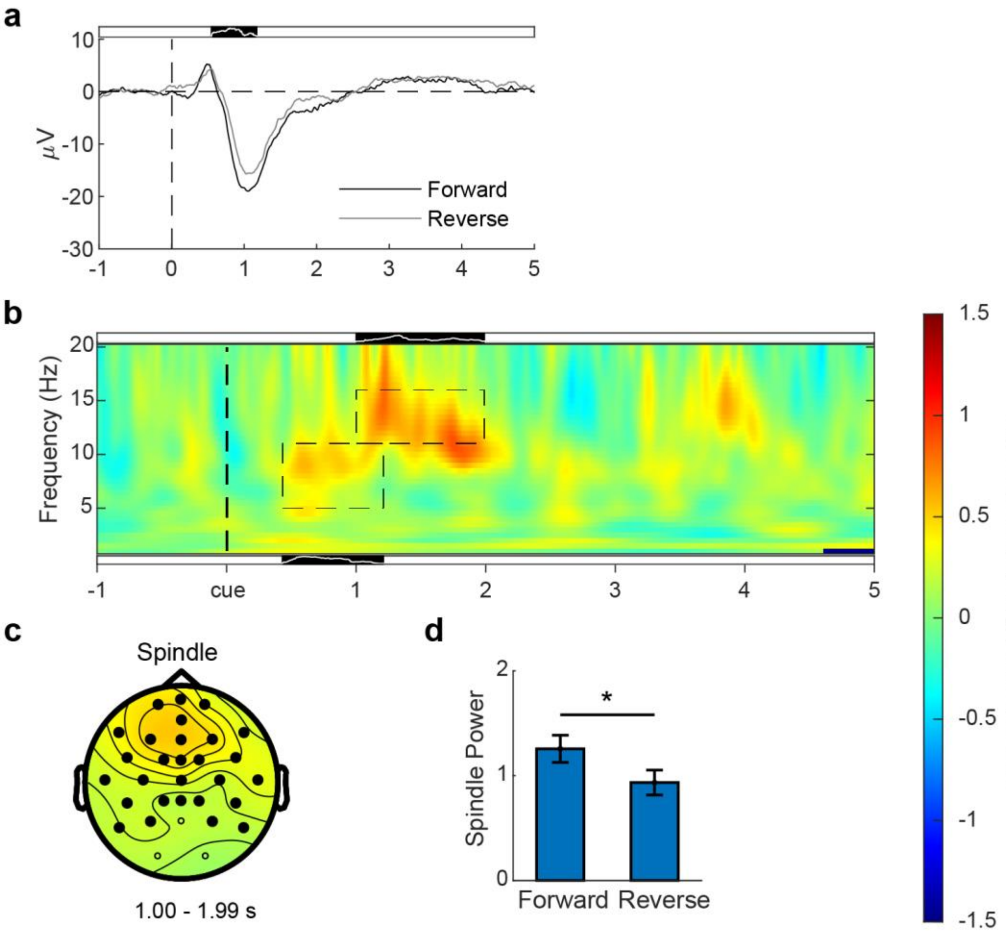


**Supplementary Fig. 2. Event-related responses of forward and reverse words.** (**a**, **b**, **c**, **d**): Presentation of understandable words during non-rapid eye movement sleep (forward words) elicited a higher negative evoked potential 548–1174 ms after cue onset compared with reverse words (cluster $P = .010$; **a**), and a higher increase in spindle power (11–16 Hz; 1000–1990 ms; cluster $P = .030$; **b**) and theta to alpha power (5–11 Hz; 430–1210 ms; cluster $P = .030$; **b**). The black areas above the plots indicate significant time intervals, while the white line in the black areas display the number of significant areas (full height = 31 electrodes). (**c**): The increase in spindle power was most prominent over frontal brain regions (significant electrodes = filled black dots). (**d**): Mean power in the significant cluster, averaged over significant electrodes, duration and frequency band of interest, was higher for forward vs. reverse words ($t(99) = 2.54$, $P = .013$). Values in (**d**) are displayed as mean ± SEM. * $P < .05$.

### Supplementary tables

**Supplementary Table 1. Sleep paramters in the night with control and relaxing words.** Means in minutes ± SEM and % ± SEM relative to sleep period time. Sleep period time (SPT) including time spent awake after sleep onset (WASO), sleep onset latency (SOL, first N1 followed by a N2 epoch), sleep stage N1 and N2, slow-wave sleep (SWS), rapid eye movement (REM) sleep and sleep efficiency (time asleep / time in bed * 100). No significant differences between both nights ($P > .13$).

| Parameter | Control night | Relax night |
|---|---|---|
| SPT | 456.98 ± 2.96 | 460.69 ± 2.37 |
| SOL (min) | 20.32 ± 2.62 | 17.24 ± 2.07 |
| WASO (min) | 12.04 ± 1.84 | 10.32 ± 1.43 |
| N1 (min) | 29.43 ± 2.13 | 28.53 ± 1.93 |
| N2 (min) | 190.89 ± 6.01 | 189.4 ± 5.32 |
| SWS (min) | 130.99 ± 7.39 | 136.21 ± 6.68 |
| REM (min) | 93.63 ± 2.62 | 96.23 ± 2.80 |
| WASO (%) | 2.67 ± 0.42 | 2.26 ± 0.32 |
| N1 (%) | 6.49 ± 0.48 | 6.21 ± 0.43 |
| N2 (%) | 41.81 ± 1.32 | 41.19 ± 1.20 |
| SWS (%) | 28.59 ± 1.58 | 29.49 ± 1.42 |
| REM (%) | 20.44 ± 0.52 | 20.84 ± 0.56 |
| SWS lat | 15.25 ± 1.10 | 14.38 ± 1.01 |
| REM lat | 81.74 ± 3.33 | 82.98 ± 3.49 |
| Sleep efficiency | 92.78 ± .79 | 93.90 ± 0.62 |

**Supplementary Table 2. Task performance in the control and relax cueing night.** Means of parameters ± SEM of the word pair associate learning task (PAL), the Regensburger Wortflüssigkeitstest (RWT), semantic verbal fluency test (SVF) and a psychomotor vigilance test (PVT). MeanRRT was calculated by dividing the RT (ms) by 1000 and then transform it reciprocally (1/RT), mean reaction time (MeanRT), number of lapses (lapses), number of false starts (false starts). Right column displays *P*-values of paired Student's *t*-tests.

| Parameter | Control night | Relax night | *P*-values |
|---|---|---|---|
| PAL presleep performance (%) | 73.13 ± 2.40 | 74.44 ± 2.14 | .30 |
| PAL postsleep performance (%) | 71.94 ± 2.56 | 71.56 ± 2.44 | .67 |
| PAL improvement (presleep = 100%) | 98.10 ± 1.06 | 95.66 ± 1.33 | .12 |
| RWT | -1.38 ± 0.85 | -0.26 ± 0.84 | .35 |
| RWT improvement (presleep = 100%) | 104.32 ± 7.18 | 104.87 ± 6.16 | .95 |
| SVF | -0.24 ± 1.27 | -1.31 ± 1.23 | .57 |
| SVF improvement (presleep = 100%) | 105.08 ± 6.43 | 108.65 ± 9.43 | .77 |
| PVT MeanRT (ms) | 337.29 ± 4.59 | 336.12 ± 4.06 | .70 |
| PVT MeanRRT | 3.05 ± 0.04 | 3.05 ± 0.36 | .86 |
| PVT lapses | 4.19 ± 0.96 | 4.56 ± 0.68 | .69 |
| PVT false starts | 6.46 ± 2.29 | 10.25 ± 5.25 | .52 |

**Supplementary Table 3. List of words played during non-rapid eye movement sleep (English translation in parentheses).**

| Relax words | Control words | Reverse words |
|---|---|---|
| abtauchen (to submerge) | anreichern (to enrich) | ausruhen (to relax) |
| angenehm (pleasant) | ausscheiden (to discard) | Blei (lead) |
| Atemzug (breath) | beansprucht (claimed) | Delphin (dolphin) |
| ausruhen (to relax) | Beispiel (example) | einfach (easy) |
| beruhigend (reassuring) | besteht (exists) | erzeugen (to produce) |
| dahintreiben (to drift) | Bildung (formation) | Magmen (magmas) |
| Delphin (dolphin) | Blei (lead) | Meer (sea) |
| einfach (easy) | Boden (soil) | Phase (phase) |
| einsinken (to subside) | eingeteilt (rationed) | |
| eintauchen (to plunge) | entstehen (to produce) | |
| entfernen (to depart) | erstrecken (to extend) | |
| entspannen (to ease) | erzeugen (to generate) | |
| entspannt (relaxed) | fest (solid) | |
| erholen (to recuperate) | Gänge (veins) | |
| Fisch (fish) | gebunden (bound) | |
| Fische (fishes) | Gesteine (rocks) | |
| geniessen (to enjoy) | Gold (gold) | |
| Korallen (coral) | Kalke (chalk) | |
| Korallenfelder (coral fields) | Kristallisation (crystallisation) | |
| langsam (slow) | Kupfer (copper) | |
| leicht (light) | Lagerstätten (deposits) | |
| loslassen (to release) | Lösungen (solutions) | |
| Meer (sea) | Magmen (magmas) | |
| müder (more tired) | Metalle (metals) | |
| mühelos (effortless) | Oberfläche (surface) | |
| Ruhe (quiet) | Öl (oil) | |
| Schlaf (sleep) | Phase (phase) | |
| schlafen (to sleep) | Prozess (process) | |
| Schlaftiefe (sleep depth) | Restschmelze (residual melt) | |

| | |
|---|---|
| Schwimmen (to swim) | Rohstoffe (resources) |
| sicher (safe) | schichtförmig (layered) |
| sinken (to sink) | Schichtung (stratification) |
| spüren (to sense) | sieden (to seethe) |
| tauchen (to dive) | stark (strong) |
| tief (deep) | Stockwerk (storey) |
| tiefer (deeper) | Stoffe (materials) |
| wahrnehmen (to perceive) | Trennung (segregation) |
| Wasser (water) | verengen (to straiten) |
| weiter (further) | willkürlich (arbitrary) |
| wohltuend (soothing) | Zinn (tin) |

**Supplementary Table 4. Cue characteristics.** All values are means ± SD of all presented words with maximum and minimum value in parentheses. Sound pressure level was measured directly at the loudspeaker.

| Parameter | Relaxing words | Control words | Reverse relaxing words | Reverse control words |
|---|---|---|---|---|
| Mean duration (s) | 1.24 ± 0.21 [0.89 – 1.67] | 0.83 ± 0.15 [0.55 – 1.20] | 1.11 ± 0.23 [0.97 – 1.45] | 0.80 ± 0.08 [0.69 – 0.88] |
| Sound pressure level (dB) | 51.00 ± 3.09 [44.80 – 57.40] | 51.61 ± 2.51 [43.70 – 55.30] | 52.43 ± 2.39 [50.30 – 55.60] | 51.08 ± 1.83 [49.00 – 53.10] |

### Supplementary Movie

**Supplementary Movie 1.** Event-related slow wave activity power of relaxing compared with control words.
